# Noradrenergic inputs to the basolateral amygdala have bidirectional effects on coping behavior and neuronal activity in mice

**DOI:** 10.1101/2025.09.18.677150

**Authors:** Alexa R. Soares, Caroline Fai, Yann S. Mineur, Marina R. Picciotto

## Abstract

Norepinephrine (NE) signaling is disrupted in stress disorders, with insufficient NE signaling implicated in major depressive disorder and hyperactive NE signaling associated with post-traumatic stress disorder, suggesting that adequate mood regulation requires optimal NE levels. The basolateral amygdala (BLA) is a hub for stress processing and receives dense noradrenergic innervation from the locus coeruleus (LC), the primary noradrenergic nucleus in the brain. The relationship between LC activity and cognitive/behavioral function during fear conditioning has been described as an inverted U, in which moderate LC activity, and subsequent NE release, is required for adaptive coping to threats, while hyperactive LC-NE signaling drives maladaptive behavioral responses. We used fiber photometry to measure NE signaling in the mouse BLA during acute behavioral responses to escapable and inescapable stressors, and then used an optogenetic approach to stimulate the noradrenergic terminals in the BLA at different frequencies to evaluate effects on coping behavior and cFos expression in the LC-BLA circuit. We found that low-frequency stimulation of the circuit inhibited both passive coping and BLA neuronal activity, while high-frequency stimulation had the opposite effect; the behavioral effects were not mediated by sex, but the cFos effects were specific to males. This study represents an expansion of the inverted U framework to encompass LC-BLA signaling driving acute behavioral responses to stress.

**Graphical abstract:** 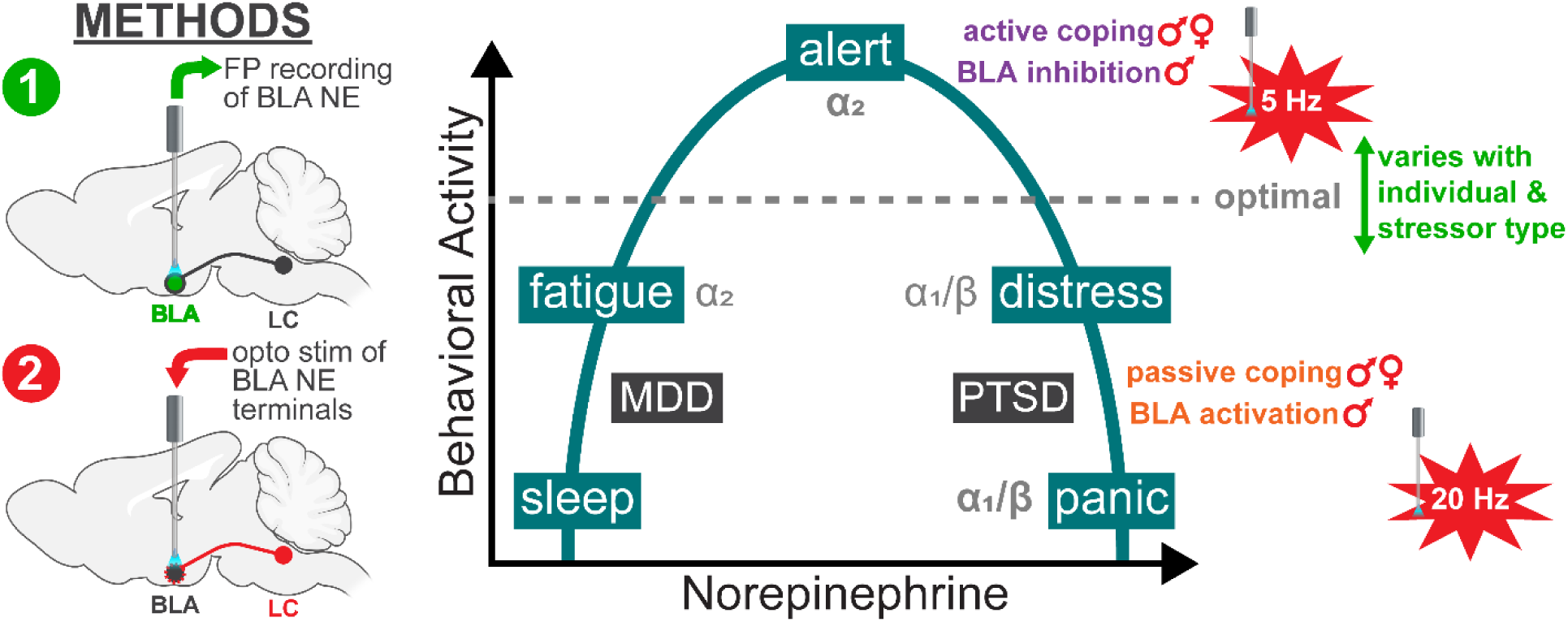

**Highlights:**

- BLA NE levels signal behavioral transitions between coping responses to stress
- Low-frequency stimulation of BLA NE terminals inhibits passive coping and BLA cFos
- High-frequency stimulation of BLA NE terminals promotes passive coping and BLA cFos
- Behavioral effects are similar across sexes, but cFos alterations are male- specific

## 1. Introduction

Disorders of stress signaling like major depressive disorder (MDD) and post-traumatic stress disorder (PTSD) represent a serious public health challenge, affecting an estimated 29.9% and 10.1% of Americans, respectively (Kessler et al., 2012). Both disorders are marked by a transition from adaptive to maladaptive coping strategies in response to stressful events (Aldao et al., 2010; Compas et al., 2017). The catecholamine norepinephrine (NE) is a well-documented neuromodulator of coping behavior (Hermans et al., 2025). Furthermore, seemingly opposite disruptions to the noradrenergic system are implicated in both MDD and PTSD: MDD is characterized by insufficient NE signaling (Hamon and Blier, 2013), whereas elevated NE is a hallmark of PTSD (Southwick, 1993). These patterns suggest that a careful balance of NE signaling is crucial to engage adaptive coping responses to stress.

Adaptive coping behavior is further hindered by the perception of uncontrollability, as the loss of control promotes continued maladaptive responses (Hermans et al., 2025; Moscarello and Hartley, 2017). This trend is heightened in individuals with stress disorders, as people with MDD (Diener et al., 2009) and PTSD (Hancock and Bryant, 2020) are more inclined towards maladaptive coping behaviors following a perceived loss of control, and persistent beliefs surrounding uncontrollability of stress are implicated in the maintenance of PTSD symptomology (Ehlers and Clark, 2000). As it is more difficult to assess what is adaptive for a rodent, preclinical models of these disorders instead refer to proactive/active, vs. reactive/passive, coping responses to stress (de Boer et al., 2017; de Kloet and Molendijk, 2016; Koolhaas et al., 1999).

Preclinical work has identified a circuit connecting cortical, limbic, and brainstem regions that coordinates defensive behaviors in response to stressful stimuli, and the amygdala has been highlighted as a critical node within this circuit (Hermans et al., 2025; Iqbal et al., 2023; Moscarello and Hartley, 2017). The amygdala integrates sensory information about the stressor determining the controllability or escapability of a threat: when high control is perceived, a pathway emanating from the amygdala to initiate active coping responses, such as escape, is engaged; under conditions of low controllability, the amygdala engages a different pathway that initiates passive coping behaviors such as freezing (Moscarello and Hartley, 2017). Several assays have been developed to explore acute responses to escapable and inescapable stressors in rodents. For example, the looming shadow test (LST), representative of an escapable stressor, provokes innate defensive behaviors to an ethologically relevant stimulus (Yilmaz and Meister, 2013). The tail suspension test (TST), representative of an inescapable stressor, also examines innate defensive responses and is a reliable predictor of antidepressant efficacy (Cryan et al., 2005).

Preclinical work has shown that NE signaling through the basolateral nucleus of the amygdala (BLA), the primary input area of the amygdala (Correia and Goosens, 2016; LeDoux and Daw, 2018), regulates behavioral responses to stress (Mineur et al., 2018; Rajbhandari et al., 2015). The BLA receives dense noradrenergic innervation from the locus coeruleus (LC; Daviu et al., 2019; de Boer et al., 2017), the main noradrenergic nucleus in the brain (Aston-Jones and Cohen, 2005; Valentino and Van Bockstaele, 2008). These LC-BLA noradrenergic projections control active and passive coping responses to acute stressors (Giustino et al., 2020; McCall et al., 2017).

The relationship between LC neuronal activity and behavioral responses, particularly in the context of fear conditioning, has been described as an inverted-U (Arnsten, 2009; Bouras et al., 2023). At baseline, LC activity is low, and thus arousal is also low; salient sensory stimuli can trigger increased firing, which increases arousal and promotes adaptive responding to the stimulus by elevating cortical control over the amygdala; with increasing stress, LC activity increases, so arousal is elevated but appropriate responsiveness to stimuli is disrupted as the amygdala becomes disinhibited (Arnsten, 2009; Aston-Jones and Cohen, 2005; Bouras et al., 2023; Valentino and Van Bockstaele, 2008). The goal of the current study was to determine how noradrenergic activity in the LC-BLA circuit controls acute behavioral coping responses to escapable and inescapable stressors (Liu et al., 2025; Mineur et al., 2022). We first measured real-time NE signaling in the BLA during exposure to the LST and TST using fiber photometry, and determined how these dynamics change at transitions between coping strategies. We then used an optogenetic approach to stimulate NE terminals in the BLA at different frequencies to reveal the relationship between LC-BLA signaling and coping behaviors during stress.

## 2. Methods

### 2.1. Animal husbandry

All procedures were approved by the Yale University Institutional Animal Care and Use Committee in compliance with the National Institute of Health’s Guide for the Care and Use of Laboratory Animals. All mice were maintained in a temperature-controlled animal facility on a 12 hr light/dark cycle (lights on at 7:00 AM). Mice were group-housed, 2-5 per cage, and provided with *ad libitum* food and water.

### 2.2. Experiment 1

#### 2.2.1. Animals

Male and female (defined by large vs. small anogenital distance at weaning) C57BL/6J mice were obtained from The Jackson Laboratory (Bar Harbor, ME) at 8-10 weeks of age.

#### 2.2.2. Surgical procedures

Mice were anesthetized using isoflurane (induced at 4% and maintained at 1.5-2%) and secured in a stereotaxic apparatus (David Kopf Instruments, Tujunga, CA). 0.5 µL of the GPCR-Activation-Based NE (GRAB-NE) sensor, AAV9-hsyn-NE1.0 (Feng et al., 2019; WZ Biosciences), was delivered unilaterally to the BLA (A/P -1.0 mm, M/L -3.0 mm, D/V -5.3 mm, relative to bregma; see Fig. 1E for diagram) at a rate of 0.1 µL/min using a 2 µL syringe with a 30 gauge, flat-tip needle (Hamilton Neuros, Reno, NV). The needle was left in place for 6 min after the infusion. A fiber-optic cannula (Doric Lenses, Quebec City, Quebec, Canada) was implanted at the infusion site using the same coordinates and affixed to the skull using opaque dental cement (3M, MD). Mice were allowed to recover in a cage with a microwavable heating pad underneath before returning to their home cage. Mice remained in their home cage for at least 3 weeks before beginning behavioral testing.

**Figure 1.**
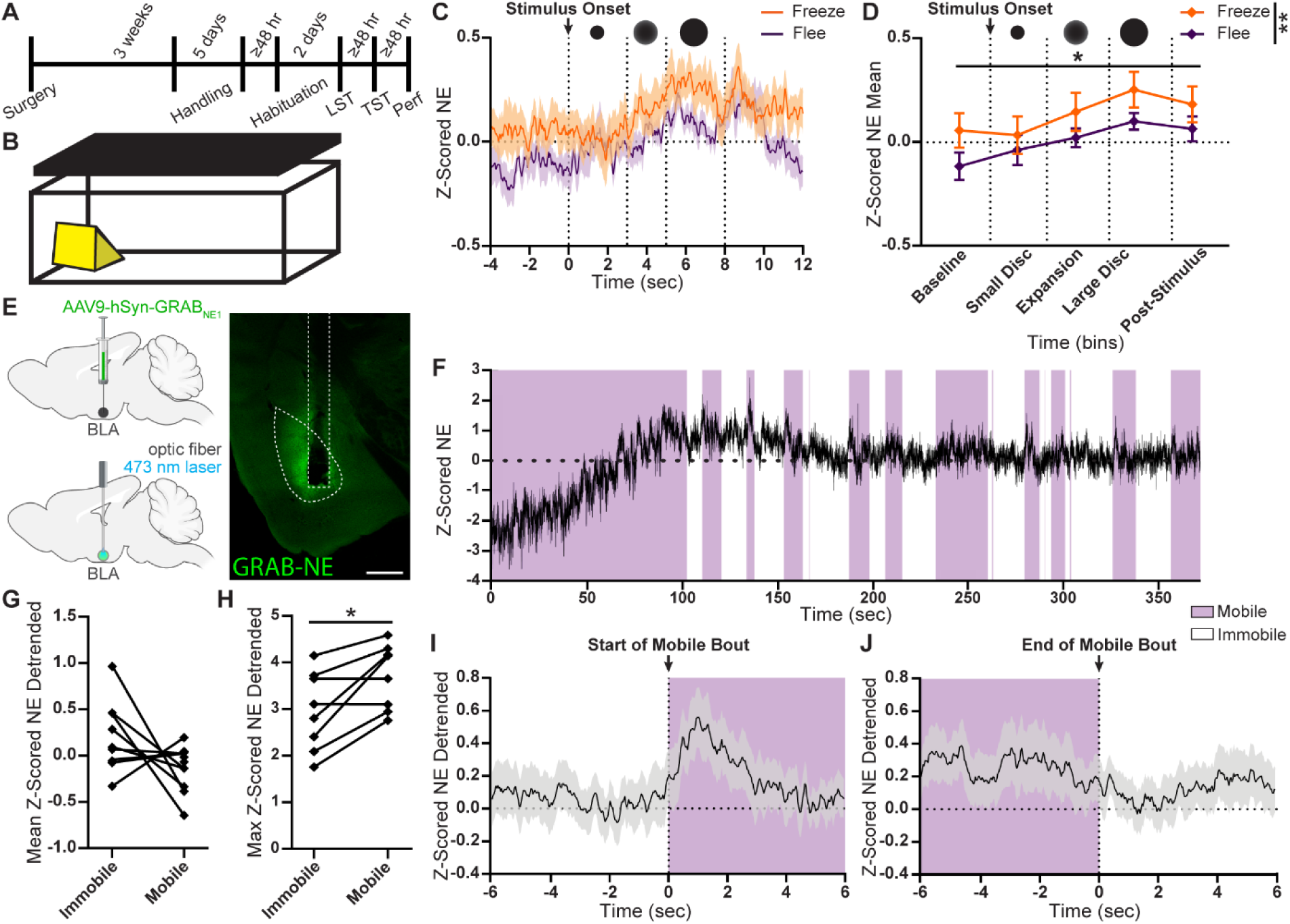
Measuring NE levels in BLA during escapable and inescapable stress. **A.** Experimental timeline. Perf = Perfusion. **B.** Diagram of LST apparatus, with shelter (yellow) in arena (white) and screen (black) above. **C.** Z-Scored fluorescence intensity corresponding to NE levels (mean ± SEM) during stimulus presentation. **D.** Mean NE fluorescence intensity (mean ± SEM), binned by stimulus phase (*n* = 10 mice). Orange indicates passive coping (freezing); purple indicates active coping (fleeing). Black circles represent stimulus phase. **E.** Top left: Diagram of AAV-GRAB-NE infusion in BLA. Bottom left: Diagram of fiber implant in BLA. Right: Representative micrograph of GRAB-NE expression and fiber tract in BLA coronal slice; scale = 500 μm. **F.** Z-scored NE fluorescence intensity during the TST for an example mouse. Purple background indicates active (mobile) coping; white background indicates passive (immobile) coping. **G.** Mean (*n* = 9 mice) and **H.** maximum Z-scored, detrended NE fluorescence intensity during active vs. passive coping (*n* = 8 mice). **I.** Z-scored NE fluorescence intensity (mean ± SEM) for an example mouse during initiation of active coping, and **J.** initiation of passive coping (*n* = 14 bouts). Purple background indicates active coping; white background indicates passive coping. \**p* < 0.05; \*\**p* < 0.01. Partially created with BioRender.com

#### 2.2.3. Fiber photometry (FP)

Fluorescent measurements of GRAB-NE sensor activity were recorded using a Doric Lenses 1-Site Fiber Photometry System. The Fiber Photometry Console controlled two connectorized LEDs (CLEDs) through the LED module driver. The isosbestic channel was stimulated through a 405 nm CLED modulated at 208.616 Hz and the green channel was stimulated through a 465 nm CLED modulated at 572.205 Hz. Each CLED was connected via attenuating patch cord to the five-port Fluorescence Minicube. A pigtailed fiber-optic rotary joint was connected to the Minicube and suspended above the behavioral chamber with a rotary joint holder to deliver and record light. The other end of the rotary joint was connected to the mono fiber-optic patch cord and attached to the implanted fiber-optic cannula on the mouse via a zirconia sleeve. The F1 output port on the Minicube was connected to the Photoreceiver (New Focus, San Jose, CA) that transmitted the 560 nm fluorescence from the GRAB-NE sensor to the Fiber Photometry Console via an analog port. The fluorescent signal was recorded using Doric Neuroscience Studio (Version 5) via the Lock-In demodulation mode with a sampling rate of 12.0 kS/sec. Data were decimated by a factor of 100. For behavioral assays requiring video recordings, a Doric USB3.0 Behavior Tracking Camera was connected to the Fiber Photometry Console via a digital port and recorded video at 30 frames/sec. For behavioral assays requiring manual timestamps of events, a custom TTL trigger was acquired from Doric and connected to the Fiber Photometry console via a digital port. Raw data from Doric Neuroscience Studio were pre-processed using custom MATLAB (R2022a, MathWorks) scripts. The first second of collected data was removed to avoid bleaching artifacts, and the data were then downsampled to match the frame rate of the camera.

To validate expression, activity, and targeting of the GRAB-NE sensor after recovery from viral infusion surgery but before handling, fluorescent signals were recorded during exposure to a mild foot shock. The optic fiber was connected to the FP system and the mouse was placed in a novel empty cage for 5 min to acclimate. The mouse was then transferred to a chamber with a grid floor (MedAssociates, VT). After 10 sec, a 1-sec, 0.4 mA shock was administered and corresponding TTL timestamp recorded. 5 sec after shock delivery, the mouse was returned to an empty cage and disconnected from the FP system. The green (= NE) channel was compared to the isosbestic channel to confirm a strong signal-to-noise ratio; mice with noisy signals were excluded from further experiments. After signal testing, mice were handled daily for 5 days before further behavioral testing.

#### 2.2.4. LST

At least 48 hr after the last day of handling (see Fig. 1A for timeline), mice underwent the LST to assess coping responses to escapable stress using a protocol based on published experiments in mice (Daviu et al., 2020a). The LST was conducted in a 24 x 15.5 x 20 cm plastic arena. A monitor the size of the arena was mounted above the arena, facing down, displaying a blank white screen. Experiments were conducted in a dark room so that the only light present was emitted from the monitor. A custom 3D-printed translucent shelter was placed in the center of one of the short sides of the arena (see Fig. 1B for diagram). The FP rotary joint was mounted such that the patch cord fit in between the monitor and the arena, and the Doric Camera was mounted on the other end of the arena to record video. For 2 days, the FP patch cord was attached, and each mouse was allowed to freely explore the arena for 20 min in order to habituate. On the third day, the LST was performed. The FP system was connected, the recording started, and the mouse was allowed to explore the arena for 3 min undisturbed. After 3 min, the 8-sec looming shadow stimulus was presented on the monitor to mimic a descending aerial predator: a small black disc appeared in the center of the screen for 3 sec (Small Disc phase), the disc then expanded for 2 sec (Expansion phase), and then remained static at its full size for 3 sec (Large Disc phase). This stimulus was presented 5 times for each mouse, with at least 1 min in between trials, and the TTL trigger was used to indicate when the stimulus was presented. A faint mark on the floor of the arena indicated the midline, and the stimulus was only presented when the mouse was across the midline from the shelter. The recording ended 1 min after the final trial. If the mouse remained inside the shelter for more than 10 min, the test was terminated.

Videos were manually scored to determine the timing and category of the coping response. Trials in which the mouse froze during stimulus presentation were defined as passive coping trials, and trials in which the mouse fled towards the shelter were defined as active coping trials. Trials in which the mouse did not exhibit a behavioral response to the stimulus were excluded from analysis.

Raw data were pre-processed as described in Section 2.2.3, and then FP data in both channels were Z-scored and the isosbestic signal was subtracted from the NE signal to correct for artifacts. For Stimulus Onset analyses, FP data were extracted for each analyzed trial starting 4 sec before the onset of the Small Disc phase (Baseline phase) through 4 sec after the end of the Large Disc phase (Post-Stimulus phase) and time- locked to the onset of the Small Disc phase. For each mouse, the data were sorted by response type and then averaged within each response category. Within each category, the average trace from each mouse was then averaged and plotted. To evaluate differences between response categories, the average FP signal for each mouse was determined for each phase of the stimulus. 3-way Repeated-Measures Analyses of Variance (rmANOVA) were performed with Time as a within-subject factor and Coping Response and Sex as between-subject factors. As significant sex differences were not observed (see Fig. S1A and Table S1), data were collapsed across Sex and 2-way rmANOVAs were performed with Time as a within-subject factor and Coping Response as a between-subject factor. Statistical analyses were performed in GraphPad Prism 10 and Microsoft Excel. To ensure visibility on graphs (Fig. 1C), traces were smoothened using a moving average with a binning factor of 5.

#### 2.2.5. TST

At least 48 hr after the LST (see Fig. 1A for timeline), mice underwent the TST to assess coping responses to an inescapable stressor (Cryan et al., 2005). Mice were connected to the FP system and immediately suspended gently by the tail and fluorescent signals were recorded for 6 min.

Raw data were pre-processed as described in Section 2.2.3, and then videos were analyzed using DeepLabCut (Version 2) for bodypart tracking (Mathis et al., 2018; Nath et al., 2019). Specifically, we labeled 8 frames/video, and 95% were used for training.

We used a ResNet-50-based neural network with default parameters for 200,000 training iterations. We validated with 1 shuffle, and found the test error was 7.81 pixels. We then used a p-cutoff of 0.6 to condition the X,Y coordinates for future analysis. This network was then used to analyze each frame in every video. Following DeepLabCut analysis, the bodypart tracking data were processed using a custom MATLAB script to calculate displacement of the base of the tail between each frame as a measure of mobility. Displacement less than 2.5 pixels/sec was defined as immobility, or passive coping, as this threshold corresponded to frames where the mouse displayed no movement except for respiration; above this threshold was defined as mobility, or active coping.

Both channels of FP data were Z-scored, and then the isosbestic signal was subtracted from the green signal to correct for movement and other signal artifacts. We observed consistent positive trend in the signal across each trial (Fig. 1F), so we linearly detrended the data for further analyses to more accurately assess differences between active and passive coping. FP data were then aligned to the transitions between active and passive coping determined by the bodypart tracking. FP data were extracted from 6 sec before each transition between coping responses through 6 sec after each transition to visualize the dynamics during these transitions. The average and maximum signal were calculated across the entire trial during active and passive coping. 2-way rmANOVAs were performed with Coping Response as a within-subject factor and Sex as a between-subject factor to identify any sex differences. As significant sex differences were not observed (see Fig. S1B-C and Table S1), data were collapsed across Sex and paired t-tests were performed to evaluate differences between coping responses. Effect sizes are presented as Cohen’s d (*d*). Statistical analyses were performed in GraphPad Prism 10 and Microsoft Excel. To ensure visibility on graphs (Fig. 1I-J), traces were smoothened with a binning factor of 5.

#### 2.2.6. Expression and placement validation

At least 48 hr after the TST (see Fig. 1A for timeline), mice were anesthetized with pentobarbital (Fatal-Plus, Vortech Pharmaceuticals) and intracardially perfused with ∼50 mL of 4% paraformaldehyde (PFA; Electron Microscopy Sciences) in phosphate-buffered saline (PBS; Gibco). Brains were extracted and post-fixed for 1 day in 4% PFA at 4°C before transferring to 30% sucrose (Millipore Sigma) in PBS at 4°C until sectioning. Brains were sectioned on a freezing microtome (Leica) at 40 μm and slices were stored in 0.02% sodium azide (Millipore Sigma) PBS solution at 4°C until mounting. Sections containing the BLA between bregma -0.59 mm and bregma -2.03 mm (Paxinos and Franklin, 2019) were mounted on slides and coverslipped with ProLong Gold antifade mounting medium (Invitrogen). To confirm appropriate expression of GRAB-NE and placement of the fiber in the BLA, images were acquired at 10x magnification using a FLUOVIEW FV10i confocal microscope to detect the green fluorophore (Olympus; see Fig. 1E for representative micrograph). Mice were excluded from analyses if green fluorescence and/or fiber tips were not observed in the BLA.

### 2.3. Experiment 2

#### 2.3.1. Animals

Male and female (defined by large vs. small anogenital distance at weaning) DBH-Cre (B6.Cg-Dbht^m3.2(cre)Pjen^/J; strain #033951) mice from The Jackson Laboratory (Bar Harbor, ME) were bred in-house, and backcrossed onto the wildtype C57BL6/J background. Lines were maintained as Cre knock-in heterozygotes and mice for this experiment were derived from heterozygote x wildtype pairings.

#### 2.3.2. Surgical procedures

Surgeries were performed as described in Section 2.2.2, but 0.5 µL of a Cre-dependent, retrograde virus containing a tdTomato fluorophore was delivered unilaterally to the BLA of DBH-Cre mice, so that the virus would be retrogradely transported to DBH^+^ cell bodies projecting to the BLA and expressed on those projections. oChIEF animals received rg-DIO-oChIEF-tdTomato (SignaGen Laboratories), expressing the blue light-activated ion channel oChIEF (Lin et al., 2009) and the tdTomato fluorophore, while tdTomato control animals received rg-DIO-tdTomato (SignaGen Laboratories) to validate viral expression and targeting. Within each cohort, all animals received the same light stimulation through the fiber implant over the BLA (see Fig. 2A for diagram). After surgery, mice were returned to their home cage for at least 2 months to ensure retrograde transport and expression of the oCHIEF construct and then handled daily for 5 days before beginning behavioral testing.

**Figure 2.**
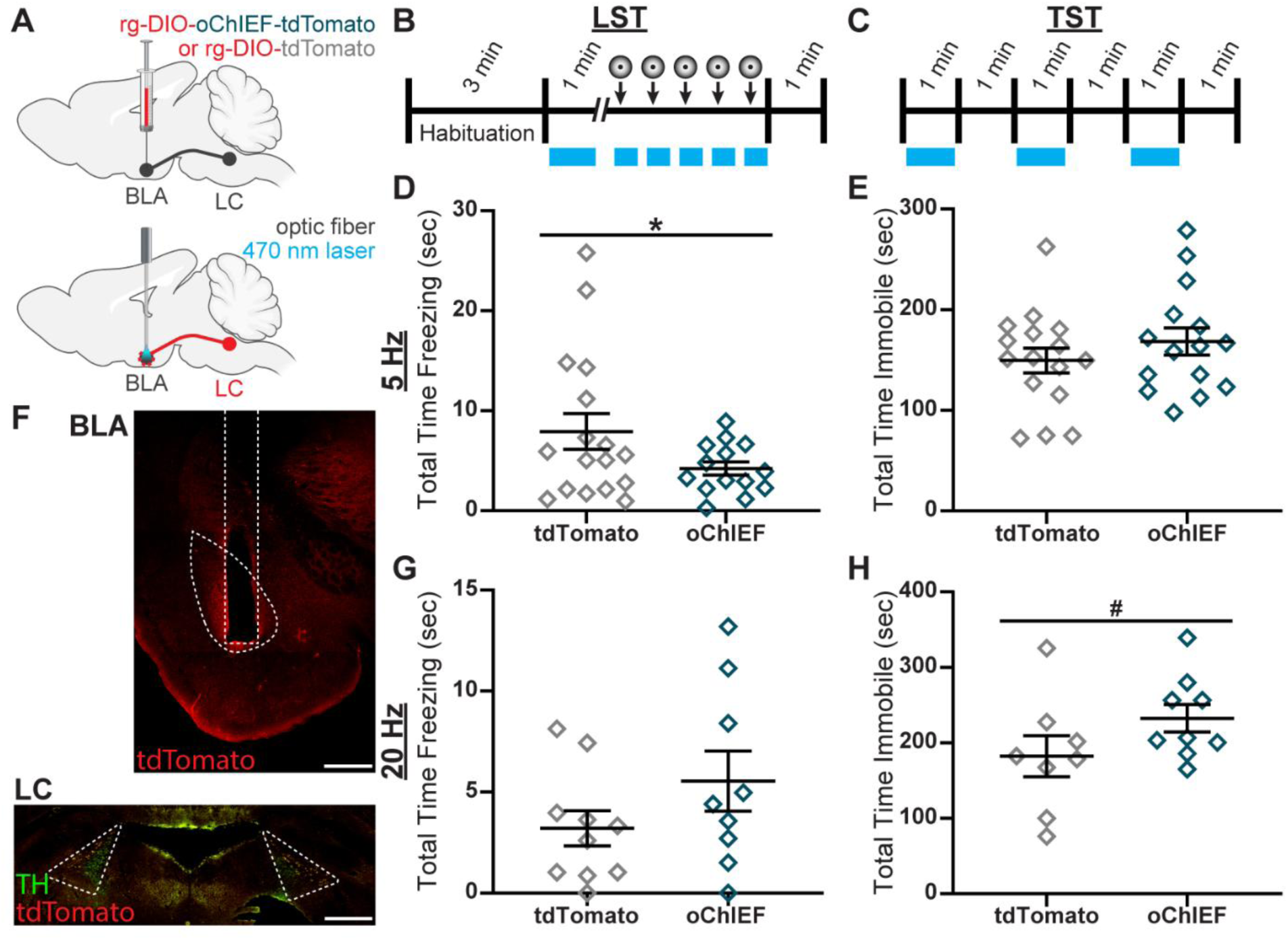
Behavioral effects of optogenetic stimulation of NE terminals in BLA. **A.** Top: Diagram of viral infusion in BLA. Bottom: Diagram of fiber implant in BLA. **B.** Timeline of stimulation during LST; circles represent looming shadow stimuli presentations. **C.** Timeline of stimulation during TST. Blue bars represent duration of laser stimulation. **D.** Effects (mean ± SEM) of 5 Hz stimulation on passive coping during LST (*n* = 14-17 mice/group) and **E.** TST (*n* = 15-16 mice/group). **F.** Top: Representative micrograph of fiber tract in BLA coronal slice; scale = 500 μm. Bottom: Representative micrograph of tdTomato expression in TH^+^ cells in LC coronal slice; scale = 500 μm; green = TH; red = tdTomato. **G.** Effects (mean ± SEM) of 20 Hz stimulation on passive coping during LST (*n* = 9-10 mice/group) and **H.** TST (*n* = 8-9 mice/group). ^#^*p* < 0.07; \**p* < 0.05. Partially created with BioRender.com

#### 2.3.3. Optogenetics

Optogenetic stimulation of BLA noradrenergic terminals was carried out using a modified version of the Doric Lenses 1-Site Fiber Photometry System described in Section 2.2.3. The CLEDs were replaced with a DPSS Blue 473 nm Laser (OEM Laser Systems, Draper, UT) driven by a PSU-III LED Power Supply (OEM Laser Systems, Draper, UT) which was connected via non-attenuating patch cord to a custom four-port Fluorescence Minicube. Separate cohorts of animals received either 5 or 20 Hz pulses, controlled by Doric Neuroscience Studio (Version 5) and delivered via BNC cable connecting the Fiber Photometry Console to the laser Power Supply. As in Section 2.2.3, the Minicube was connected to the cannula via a rotary joint with a mono fiber-optic patch cord and zirconia sleeve, and the Doric USB3.0 Camera was used to record synchronized video. The laser was powered to provide an output of 10 mW at the tip of the zirconia sleeve, as measured by a Photodiode Sensor (ThorLabs, Dachau, Germany).

#### 2.3.4. Open field test (OFT)

At least 48 hr after handling, mice underwent the OFT to determine effects of the optogenetic stimulation on baseline locomotion. After being connected to the patch cord, mice were placed in a 38 x 38 x 13 cm plastic arena for 6 min. 5 or 20 Hz laser stimulation was provided in 1-min intervals, starting with the laser on for the first minute.

Videos were analyzed over several cohorts using DeepLabCut (Version 2) for multi-animal bodypart tracking to label both the mouse and the borders of the arena (Mathis et al., 2018; Nath et al., 2019). We labeled 5-40 frames/video, and 95% were used for training. We used DLCRNet-MS5-based neural networks with default parameters for 50,000-97,000 training iterations. We validated with 1 shuffle, and found the test error was 3.64-4.86 pixels. We then used a p-cutoff of 0.6 to condition the X,Y coordinates for future analysis. These networks were then used to analyze each frame in every video.

Following DeepLabCut analysis, the bodypart tracking data were processed using a custom MATLAB script to calculate displacement of the mouse’s headcap as a measure of distance traveled. Total distance traveled across the entire trial was summed as the measure of locomotion. 2-way ANOVAs were performed with Virus and Sex as between-subject factors to evaluate sex differences. As significant sex differences were not observed (see Fig. S2 and Table S1), data were collapsed across Sex and t-tests were performed to evaluate effects of the stimulation, with Welch’s correction applied when appropriate for groups with unequal variances. Effect sizes are presented as Cohen’s d (*d*). Statistical analyses were performed in GraphPad Prism 10 and Microsoft Excel.

#### 2.3.5. LST

At least 48 hr after the OFT, mice underwent the LST as described in Section 2.2.4, with optogenetic stimulation during the test. After the initial 3-min habituation period, 5 or 20 Hz laser stimulation was provided for 1 min. At least 1 min after the end of this stimulation, the first looming stimulus was presented; laser stimulation was provided for the duration of each looming stimulus (see Fig. 2B for timeline).

As described in Section 2.2.4, videos were manually scored to determine coping responses. Total time spent freezing across all 5 trials was summed as the measure of passive coping. Statistical analyses were performed as described in Section 2.3.4.

#### 2.3.6. TST

At least 48 hr after the LST, mice underwent the TST as described in Section 2.2.5, with optogenetic stimulation during the test. 5 or 20 Hz laser stimulation was provided in 1- min intervals, starting with the laser on for the first minute (see Fig. 2C for timeline).

Videos were analyzed as described in Section 2.2.5 over several cohorts using DeepLabCut. We labeled 7-34 frames/video, and 95% were used for training. We used ResNet-50-based neural networks with default parameters for 500,000-525,000 training iterations. We validated with 1 shuffle, and found the test error was 3.00-4.01 pixels. We then used a p-cutoff of 0.6 to condition the X,Y coordinates, and used these networks to analyze each frame in every video. As described above, a custom MATLAB script was used to determine passive coping, using thresholds of 0.4-2.0 pixels/sec due to differences in the camera placement. Total time spent immobile across the entire trial was summed as the measure of passive coping. Statistical analyses were performed as described in Section 2.3.4.

2.3.7. Drugs

For the 5 Hz cohort, N-(2-chloroethyl)-N-ethyl-2-bromobenzylamine (DSP-4) was dissolved in PBS (Gibco) at 5 mg/mL and delivered at 10 mL/kg bodyweight intraperitoneally (IP) for a dose of 50 mg/kg. Mice received daily injections over 3 days, with the last injection occurring 24 hr before cFos stimulation. Control animals received 10 mL/kg bodyweight of PBS IP. As no significant effect of DSP-4 was observed (see Fig. S3A-B and Table S1), this procedure was not continued for the 20 Hz cohort.

Instead, all animals simply received 10 mL/kg bodyweight of PBS IP 30 min before the cFos stimulation.

#### 2.3.8. cFos stimulation

At least 48 hr after the TST, mice received a final laser stimulation before perfusing to examine the effects of the stimulation on cFos expression in the brain. Mice were connected to the patch cord and placed in the OFT arena for 6 min. 5 or 20 Hz laser stimulation was provided in 1-min intervals, starting with the laser on for the first minute. Mice were returned to their home cage for 90 min before perfusing to allow for maximal cFos protein expression (Chaudhuri et al., 2000). Mice were perfused and brains were post-fixed, sectioned, and stored as described in Section 2.2.6 before immunohistochemistry (IHC).

#### 2.3.9. IHC

Sections containing the BLA between bregma -0.59 mm and bregma -2.03 mm and the LC between bregma -5.33 mm and -5.99 mm (Paxinos and Franklin, 2019) were washed in PBS (Gibco) and then blocked in 3% normal donkey serum (NDS; Jackson Immuno) and 0.3% Triton-X (American Bioanalytical) PBS solution for 1 hr at room temperature (RT). Sections were then incubated in primary sheep anti-TH antibody (Sigma AB1542) at a concentration of 1:2000 and rabbit anti-cFos antibody (Cell Signaling Technologies 2250) at a concentration of 1:1000 in PBS overnight at 4°C. On day 2, sections were rinsed in PBS and incubated in secondary donkey anti-sheep 488 antibody (Abcam ab150177) and donkey anti-rabbit 647 antibody (Invitrogen A-31573), each at a concentration of 1:1000 in PBS for 1 hr at RT. Sections were rinsed again in PBS, then mounted on slides and coverslipped with ProLong Gold antifade mounting medium (Invitrogen).

#### 2.3.10 cFos analysis

Multichannel images of TH staining in the green channel, cFos staining in the far-red channel, and a 580 nm filter to detect tdTomato expression in the red channel were acquired at 10x magnification of the BLA and LC using a FLUOVIEW FV10i confocal microscope with a numerical aperture of 2.5x. At least 3 images of the BLA were captured unilaterally with the fiber tract. Acquired images were analyzed first in FIJI (Schindelin et al., 2012) with a custom Macro code to determine the area of the BLA and manually count the cFos^+^ cells. Subsequently, a custom MATLAB code was used to sum this count across all images for a given animal and divide by the total area imaged for that animal to determine density of cFos^+^ cells in the BLA. At least 3 images were captured bilaterally of the LC in each animal. Manual counting was performed to determine the number of tdTomato^+^ cells and the number of tdTomato^+^ cells co- expressing cFos. Subsequently, a custom MATLAB code was used to sum both counts across all images for a given animal and divide the number of co-expressing cells by the total tdTomato^+^ cells to determine the percentage. 2-way ANOVAs were performed with Virus and Sex as between-subject factors to identify any sex differences in the effects of the stimulation on cFos expression in both brain regions. Significant effects were followed up with *post-hoc* Šídák’s multiple comparisons t-tests when appropriate, as the between-subject factors were independent. Effect sizes are presented as partial eta squared (*η_p_*^2^) for ANOVAs and Cohen’s d (*d*) for t-tests. Simple linear regressions were performed to determine the correlation between cFos expression in the LC and BLA. Goodness of fit is presented as the coefficient of determination (*R*^2^). Statistical analyses were performed in GraphPad Prism 10 and Microsoft Excel.

#### 2.3.11. Expression and placement validation

To confirm appropriate placement of the fiber in the BLA, images were acquired at 10x magnification using a FLUOVIEW FV10i confocal microscope with a 580 nm filter to detect tdTomato (see Fig. 2F for representative micrograph). Mice were excluded from analyses if fiber tips were not observed in the BLA.

To confirm appropriate retrograde transport of the virus to TH^+^ cells in the LC, multichannel images were acquired at 10x magnification of the tdTomato fluorophore in the red channel and TH staining in the green channel using a FLUOVIEW FV10i confocal microscope (see Fig. 2F for representative micrograph). Mice were excluded from analyses if colocalization was not observed between TH^+^ and tdTomato^+^ cells in the LC.

## 3. Results

### 3.1. Elevated levels of NE in BLA precede passive coping responses to escapable stress

We have previously shown that blocking NE signaling in the BLA impairs coping responses to acute stress (Mineur et al., 2018), but the temporal dynamics of BLA NE signaling during exposure to different types of stressors remains unknown. We first set out to investigate how BLA NE dynamics change based on the type of stressor and the mouse’s coping response. Extracellular levels of NE in the BLA were measured via FP recordings of GRAB-NE fluorescence (Feng et al., 2019) in the BLA of wildtype mice (Fig. 1E) during exposure to escapable and inescapable stressors.

To examine the noradrenergic response to an escapable stressor, we recorded extracellular NE dynamics in the BLA during the LST (Fig. 1B). An active coping response was defined as fleeing towards the shelter and the passive coping response as freezing in place. When we aligned the NE dynamics to the onset of the looming stimulus, we observed that NE levels rose as the shadow expanded, potentially signaling the advancing threat, regardless of coping response; higher NE levels before the stimulus onset were associated with passive coping (Fig. 1C). Indeed, when we quantified the mean NE fluorescence during each phase of the stimulus, there was a significant effect of time (*F*4,90 = 2.664, *p* = 0.0375, Fig. 1D) and a significant difference between active and passive coping trials (*F*1,90 = 7.385, *p* = 0.0079, Fig. 1D). This pattern suggests that heightened levels of BLA NE at baseline may predispose animals to a passive coping response to inescapable stress.

### 3.2. BLA NE levels rise during active coping to inescapable stress

At least 48 hr later, we recorded extracellular NE dynamics in the same cohort of animals during the TST to examine the BLA noradrenergic response to inescapable stress. Throughout the 6-min TST, mice switch back and forth between active coping (mobile bouts) and passive coping (immobile bouts), generally starting with a prolonged mobile bout. We observed that NE levels in the BLA rise dramatically during this first bout, continue rising during each subsequent mobile bout, and never return to baseline, resulting in an overall positive trend across the duration of the trial (Fig. 1F). We linearly detrended the signal to assess differences between active and passive coping, and found a consistent spike in BLA NE immediately after transitions into active coping (Fig. 1I), with no distinctive change in NE around transitions into passive coping (Fig. 1J), highlighting that changes in the BLA NE signal are specific to active coping bouts.

Furthermore, we found that although the mean BLA NE signal doesn’t differ between active and passive coping (*t* = 1.733, *p* = 0.1214, Fig. 1G), there is a significant increase in the maximum fluorescence during active coping (*t* = 3.406, *p* = 0.0113, *d* = 1.703, Fig. 1H), which can be attributed to that spike (Fig. 1I). Thus, the relationship between BLA NE signaling and coping behavior appears to depend on the controllability of the stressor. In both assays, high levels of NE leads to subsequent passive coping. In the context of escapable stress, high baseline levels of NE may predispose the animal to passive coping, while increasing NE release generated during inescapable stress may have a similar effect.

### 3.3. Optogenetic stimulation of BLA NE terminals has frequency-dependent effects on coping behavior

Next, we used an optogenetic approach to probe the causal relationship between NE release in the BLA and coping responses to these acute stressors. We induced expression of the blue light-activated ion channel oChIEF (Lin et al., 2009) tagged with the tdTomato fluorophore in noradrenergic neurons projecting to the BLA(Fig. 2A) and provided 473-nm laser stimulation via an optic fiber over the BLA (see Fig. 2F for representative micrograph). Control animals received only the tdTomato fluorophore but received the same laser stimulation to account for any effects of the light itself (Fig. 2A). To probe the relationship between LC activity and behavior, we explored frequency-dependent effects by stimulating the noradrenergic terminals in the BLA at either 5 or 20 Hz in separate cohorts of mice.

In order to simulate the elevated NE baseline we observed in passive coping trials during the LST (Fig. 1C), we provided optogenetic stimulation of BLA NE terminals during the first min of the test, and then additional stimulation during each 8-sec looming stimulus presentation (see Fig. 2B for timeline), and observed effects on passive coping as measured by the total duration of freezing in response to the looming stimulus. We found that low-frequency stimulation induced a significant decrease in passive coping (*t* = 1.932, *p* = 0.0337, *d* = 0.6973, Fig. 2D), while high-frequency stimulation had no effect (*t* = 1.391, *p* = 0.0910, Fig. 2G).

To evaluate frequency-dependent effects of BLA NE stimulation on inescapable stress, we then administered 5 or 20 Hz blue light in 1-min intervals during the TST (see Fig. 2C for timeline), and measured passive coping as the total time spent in immobility.

There was no effect of the low-frequency stimulation (*t* = 1.025, *p* = 0.1569, Fig. 2E), but we observed an increase in immobile time following high-frequency stimulation that did not reach significance (*t* = 1.567, *p* = 0.0689, *d* = 0.7614, Fig. 2H). The same stimulation paradigms had no effects on locomotor activity in the OFT (5 Hz: *t* = 0.7318, *p* = 0.4700, Fig. S2A; 20 Hz: *t* = 1.318, *p* = 0.2060, Fig. S2E), so these findings cannot be attributed to changes in ambulation. These data reveal frequency-dependent, task-specific effects of NE projections to the BLA on passive coping behavior, suggesting that low-frequency firing associated with low NE release promotes active coping to escapable stress, while high-frequency firing, or high NE release, promotes passive coping to inescapable stress.

### 3.4. Neuronal activity in the LC and BLA are inversely correlated in male mice, and male BLA activity is differentially altered by low- and high-frequency stimulation

To probe effects of this stimulation on neuronal activity in the LC-BLA circuit, we measured cFos expression in both brain regions following 3-min optogenetic stimulation. Despite finding no significant sex differences in the behavioral assays (Fig. S2 and Table S1), we observed a consistent increase in cFos expression among tdTomato^+^ cells in the LC of females compared to males, independent of optogenetic stimulation (5 Hz: *F*_1,8_ = 21.72, *p* = 0.0016, *η_p_*^2^ = 0.7309, Fig. 3B; 20 Hz: *F*_1,25_ = 17.42, *p* = 0.0003, *η_p_*^2^ = 0.4107, Fig. 3D). When we measured cFos expression in the BLA, we observed a significant decrease in females in the low-frequency cohort (*F*1,14 = 11.13, *p* = 0.0049, *η_p_*^2^ = 0.4430, Fig. 3A), and an interaction between sex and virus group (*F*_1,14_ = 4.591, *p* = 0.0502, *η_p_*^2^ = 0.2469, Fig. 3A). *Post-hoc* analyses revealed a stimulation- induced decrease in BLA cFos expression specific to males, although this was not statistically significant (*t* = 2.441, *p* = 0.0563, *d* = 1.544, Fig. 3A). Furthermore, high- frequency stimulation induced a significant increase in BLA cFos expression (*F*1,25 = 7.878, *p* = 0.0096, *η_p_*^2^ = 0.2396, Fig. 3C), that appears to be driven by the males (*t* = 2.219, *p* = 0.0358, *d* = 1.344, Fig. 3C). These data suggest that, particularly in males, low-frequency stimulation of LC-NE terminals inhibits BLA neuronal activity, while high-frequency stimulation has the opposite effect, and these effects likely do not back- propagate to the cell bodies in the LC.

**Figure 3.**
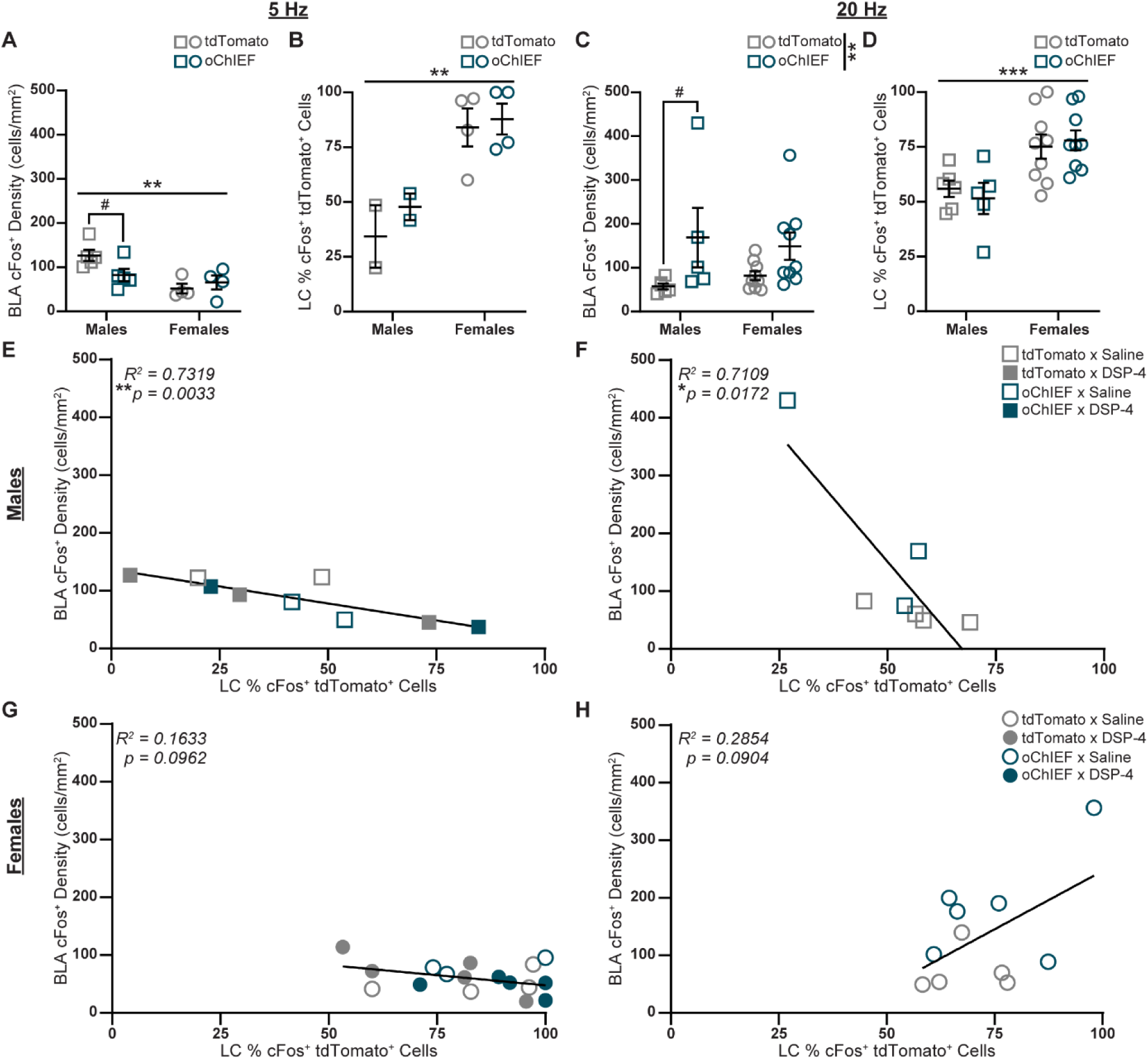
Sex-specific effects of optogenetic stimulation of BLA NE terminals on cFos levels in BLA and LC. **A.** Effects (mean ± SEM) of 5 Hz stimulation on density of cFos^+^ cells in BLA (*n* = 4-5 mice/group) and **B.** percentage of cFos^+^ cells expressing the viral fluorophore tdTomato in LC (*n* = 2-4 mice/group). **C.** Effects (mean ± SEM) of 20 Hz stimulation on BLA cFos^+^ cell density (*n* = 3-6 mice/group) and **D.** percentage of cFos^+^ tdTomato^+^ cells in LC (*n* = 3-6 mice/group). Lines indicate main effects using 2- way ANOVA; brackets indicate effects following post-hoc Šídák’s multiple comparisons tests. **E.** Linear correlation between percentage of cFos^+^ tdTomato^+^ cells in LC and cFos^+^ cell density in BLA of male mice following 5 Hz (*n* = 9 mice) or **F.** 20 Hz stimulation (*n* = 7 mice). **G.** Linear correlation between percentage of cFos^+^ tdTomato^+^ cells in LC and cFos^+^ cell density in BLA of female mice following 5 Hz (*n* = 18 mice) or **H.** 20 Hz stimulation (*n* = 11 mice). Squares indicate males, circles indicate females; open points indicate saline, closed points indicate DSP-4; gray points indicate tdTomato, teal points indicate oChIEF. ^#^*p* < 0.07; \**p* < 0.05; \*\**p* < 0.01.

We next calculated linear correlations comparing cFos expression in both brain regions to assess the relationship between LC and BLA neuronal activity. In both cohorts, we observed a significant inverse correlation between LC and BLA cFos expression, specifically in males (5 Hz: *R*^2^ = 0.7319, *F*1,7 = 19.11, *p* = 0.0033, Fig. 3E; 20 Hz: *R*^2^ = 0.2721, *F*1,20 = 7.475, *p* = 0.0128, Fig. 3F) and not females (5 Hz: *R*^2^ = 0.1633, *F*1,16 = 3.123, *p* = 0.0962, Fig. 3G; 20 Hz: *R*^2^ = 0.0003, *F*1,35 = 0.0091, *p* = 0.9245, Fig. 3H). It is important to note that some animals in the 5 Hz cohort received daily injections of DSP- 4 for 3 days preceding the final stimulation. Only the saline controls were included in these Figures 3A-B, but we did not observe an effect of DSP-4 administration on BLA (*F*1,25 = 1.125, *p* = 0.2990, Fig. S3A) or LC cFos (*F*1,19 = 0.0004, *p* = 0.9842, Fig. S3B), so we included these animals in Figures 3F-H and did not continue DSP-4 administration for the 20 Hz cohort. Together, these data suggest an inhibitory relationship between activity of pre-synaptic LC neurons and post-synaptic BLA neurons among males, and that BLA activity may be associated with passive coping.

## 4. Discussion

In the current study, we explored the role of the LC-BLA NE circuit in behavioral coping responses to escapable and inescapable stressors. Our FP data revealed task-specific patterns of NE signaling time-locked to initiation of active and passive coping, and subsequent optogenetic stimulation of NE terminals in the BLA revealed frequency-dependent effects on both coping behavior and BLA neuronal activity. Behavioral effects were task-specific, highlighting the role of escapability in mediating defensive responses. The inescapable stress assay, the TST, has been used as a reliable and high-throughput predictor of antidepressant efficacy (Cryan et al., 2005; Steru et al., 1985; Stukalin et al., 2020; Thierry et al., 1984), while the escapable stressor, the LST, was developed more recently and provides strong ethological validity of behavioral responses to a visual threat, generating behavioral and neurobiological responses that are highly conserved across species (Hu et al., 2017; Ren and Tao, 2020; Wu and Zhang, 2023; Yilmaz and Meister, 2013). Incorporating both assays allowed us to explore the role of controllability in threat processing.

Our FP data revealed that BLA NE levels rise as the shadow expands, signaling the incoming threat; similar patterns of NE reactivity to looming stimuli have been observed in the zebrafish tectum (Feng et al., 2019). Furthermore, we found that higher levels of NE, even before the onset of the stimulus, were associated with freezing responses, which may suggest that an influx of NE into the BLA signals heightened anxiety or arousal and shifts the system towards passive coping. These findings build on clinical and preclinical literature demonstrating a role for the amygdala, particularly the BLA, in responding to looming stimuli (Barbour et al., 2020; Coker-Appiah et al., 2013; Hu et al., 2017; White et al., 2018). The amygdala has previously been identified as an important node integrating sensory information from upstream areas like the superior colliculus (SC) to direct behavioral responses via projections to downstream brainstem areas (Daviu et al., 2020b; Hu et al., 2017; Huang et al., 2017; Kim et al., 2020; Laing et al., 2023; Li et al., 2018; Liu et al., 2025; Ren and Tao, 2020; Shang et al., 2018, 2015; Wei et al., 2015; Wu and Zhang, 2023; Zhou et al., 2019). Several critical pathways have been identified by which the SC relays through the BLA to drive a behavioral coping response to looming stimuli (Liu et al., 2025; Wei et al., 2015). Thus, the BLA represents an opportunity for neuromodulatory inputs, such as NE, to integrate with sensory information and shape the behavioral output (Ren and Tao, 2020).

To assess the levels of BLA NE during exposure to an inescapable stressor, we also conducted FP recordings during the TST. We have previously demonstrated a role of BLA NE, specifically signaling through α2A receptors, in mediating coping behavior during this assay (Mineur et al., 2018). Our FP data demonstrated a persistent increase in BLA NE levels across the duration of the test, with spikes in NE release overlaid during each bout of active coping, potentially representing an arousal signal. Unlike in the LST, the BLA NE signal during the TST follows, rather than precedes, coping transitions. This may highlight the role of controllability in mediating BLA threat processing. Due to other differences between these two assays, changes in NE dynamics are likely not exclusively due to perceptions of control.

We then employed an optogenetic strategy to stimulate noradrenergic terminals in the BLA at different frequencies to probe the causal relationship between BLA NE signaling and behavioral coping responses. We demonstrated bidirectional effects of this stimulation: 5 Hz stimulation of BLA NE terminals decreased passive coping to the escapable stressor, while 20 Hz stimulation increased passive coping to the inescapable stressor. In contrast, a similar study using a 5 Hz stimulation of TH^+^ LC terminals in the BLA observed an increase in anxiety-like behavior, including avoidance of the center of an OFT and the open arms of an elevated zero maze (McCall et al., 2017), which could be interpreted as increased passive coping. However, McCall and colleagues used a different opsin packaged in an anterograde viral construct, which may result in higher expression (Hui et al., 2022) and possibly greater sensitivity to light stimulation, so it is difficult to determine whether their 5 Hz stimulation is comparable to ours. Another study that directly stimulated LC neuronal cell bodies found that 25 Hz stimulation increased spontaneous freezing, while 10 Hz did not; they also found that the 10 Hz stimulation had appetitive effects in a conditioned preference test, whereas 25 Hz was aversive, and these effects were blocked by infusing α- and β-adrenergic antagonists into the BLA (Ghosh et al., 2021). These findings may represent an analogous phenomenon to the results of the present study: low-frequency LC stimulation promotes appetitive/active coping behaviors, and high-frequency LC stimulation drives aversive/passive coping behaviors, mediated by NE signaling in the BLA.

Thus, the canonical inverted-U shape of behavioral responses invoked by activity of the NE system may be expanded from a description of fear and memory processes (Bouras et al., 2023) to include acute behavioral responses to stressors, as demonstrated by the present study. In this framework, low-frequency stimulation of the LC-BLA projections represents the peak of the inverted-U in behavioral response, where moderate LC activity promotes alertness and optimal attentiveness to stressful stimuli (Arnsten, 2009), resulting in active coping responses. High-frequency stimulation simulates NE signaling toward the right side of the inverted-U-shaped response, in which high LC activity results in hyperarousal, generating a panicked, high-stress state (Bouras et al., 2023) and provoking passive coping responses. Indeed, other studies have shown that LC stimulation mimics the behavioral effects of stress on active coping in the LST (Li et al., 2018). Within this framework, we would expect low-frequency stimulation of NE terminals to inhibit BLA postsynaptic activity, and high-frequency stimulation to have the opposite effect. The cFos data suggests this may be true, at least in males: low-frequency stimulation decreased cFos expression in BLA neurons, and high-frequency stimulation increased BLA cFos, suggesting bidirectional changes in neuronal activity.

Interpretation is complicated by the sex-specificity of these effects. Despite the lack of sex differences in behavioral effects of BLA NE terminal stimulation, the cFos effects are restricted to males. Previous work from our lab found a similar pattern, in which systemic α2A agonism decreased passive coping in both males and females, while decreasing BLA cFos in males and increasing BLA cFos in females (Mineur et al., 2015). In the present study, we also observed a consistent sex difference such that LC cFos was higher in females compared to males, independent of stimulation. This may indicate a potential ceiling effect in which the female LC was already maximally activated, so optogenetic stimulation could not induce a post-synaptic effect on BLA neurons. It has been previously shown that dendrites in the female LC are longer than that of male counterparts, and in some rat strains, the female LC is larger, containing more neurons (Bangasser et al., 2016). Furthermore, estrogens have been shown to reduce expression of postsynaptic adrenergic receptors (Bangasser et al., 2016), so the female BLA may be less responsive to optogenetic stimulation of presynaptic terminals. This sexual dimorphism may explain the sex-specificity of the correlational data as well: the strong inverse correlations between male LC and BLA cFos levels suggest tighter coupling of these two regions among males. Indeed, sex hormones are known to alter functional connectivity in the human brain (Engman et al., 2016; Kogler et al., 2023), so the behavioral effects seen in females may be a result of changes elsewhere in the circuit.

Future work should investigate the post-synaptic mechanisms and downstream signals underlying the present findings. The inverted-U-shaped behavioral responses to NE signaling depends on the diversity of noradrenergic receptors: low to moderate levels of NE release activate inhibitory, high-affinity α2 receptors; as NE levels increase, the excitatory α1 and β receptors are recruited (Bouras et al., 2023). Thus, it is likely that the effects of the low-frequency stimulation are mediated by α2 receptors and the effects of the high-frequency stimulation are mediated by α1 and β receptors, but future work could employ pharmacological strategies to investigate these relationships. β receptor antagonism has been shown to block the excitatory effects of stress on BLA glutamatergic neurons (Aukema et al., 2024), and both α and β receptors in the BLA were implicated in the frequency-dependent appetitive and aversive effects of LC stimulation (Ghosh et al., 2021). We observed negative correlations between LC and BLA cFos expression in males when collapsing across groups. The mice were not exposed to stress on the day of perfusion, so this inverse relationship likely corresponds to the first two-thirds of the inverted-U-shaped response, in which LC-NE signaling through α2 receptors drives inhibition of the BLA. Another open question is the downstream pathway by which the BLA mediates behavioral responses, likely through the central nucleus of the amygdala (CeA), the primary output nucleus (Fadok et al., 2018) which has also been implicated in mediating defensive responses to looming stimuli (Ren and Tao, 2020; Shang et al., 2015; Zhou et al., 2019). In a conditioned fear paradigm, it was shown that the BLA differentially innervates competing populations of inhibitory neurons in the CeA, and this competition acts as a switch between active and passive coping responses (Fadok et al., 2017). Future work could investigate how noradrenergic inputs to the BLA modulate this switch.

Together, these experiments present an inverted-U-shaped framework for LC-BLA control of coping responses to acute escapable and inescapable stressors. Moderate levels of NE release, induced by low-frequency stimulation of NE terminals in the BLA, promote active coping to escapable stress by inhibiting BLA neuronal activity.

Conversely, high levels of NE release, induced by high-frequency stimulation of these terminals, drive passive coping to inescapable stress by activating BLA neurons. The BLA more readily facilitates a bias towards active coping when perceived control is higher, and passive coping when the stressor is uncontrollable. Coupling between LC and BLA neurons appears to be stronger in males, and future work is needed to understand the circuit mechanisms driving behavioral responses in females.

## CRediT authorship contribution statement

**Alexa R. Soares:** Conceptualization, Data curation, Formal analysis, Investigation, Methodology, Project administration, Software, Validation, Visualization, Writing – original draft. **Caroline Fai:** Formal analysis, Investigation, Validation, Writing – review & editing. **Yann S. Mineur:** Conceptualization, Funding acquisition, Methodology, Writing – review & editing. **Marina R. Picciotto:** Conceptualization, Funding acquisition, Project administration, Resources, Supervision, Writing – review & editing.

## Acknowledgements

We would like to thank Dr. Kelsea Gildawie for guidance in developing the IHC protocols, Drs. Jaideep Bains and Nuria Daviu for advice in setting up the LST, Qian Xu and Dr. Rui Chang for providing DBH-Cre mice, Samantha Sheppard for assistance with animal husbandry, and Nadia Jordan-Spasov for assistance with reagents. This paper is based upon a dissertation submitted to fulfill in part the requirements for the Degree of Doctor of Philosophy at Yale University.

