## Supplemental Material for "Noradrenergic inputs to the basolateral amygdala have bidirectional effects on coping behavior and neuronal activity in mice"

*Figure S1. Measuring NE levels in BLA during escapable and inescapable stress*

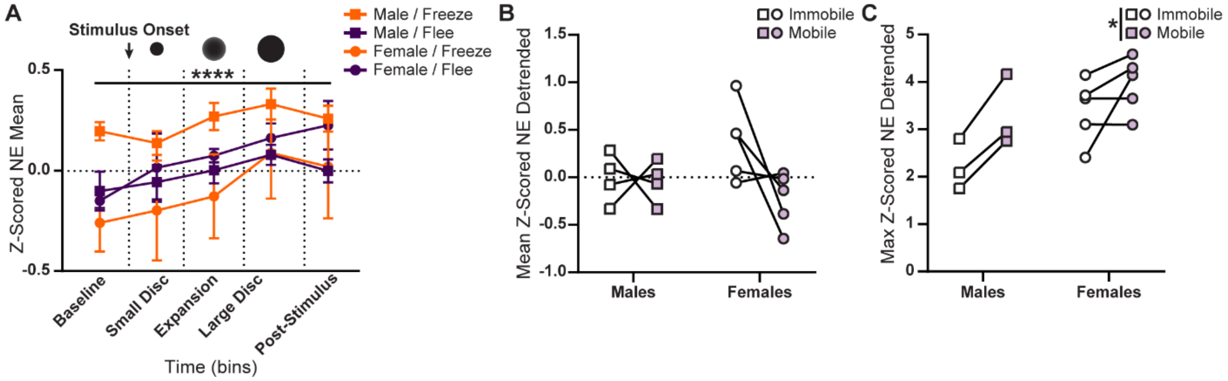


**A.** Mean NE fluorescence intensity (mean ± SEM), binned by stimulus phase in male and female mice ($n$ = 3-7 mice/sex). Orange indicates passive coping (freezing); purple indicates active coping (fleeing). Black circles represent stimulus phase. **B.** Mean ($n$ = 4-5 mice/sex) and **C.** maximum Z-scored, detrended NE fluorescence intensity during active vs. passive coping ($n$ = 3-5 mice/sex). Squares indicate males, circles indicate females. *$p < 0.05$; ****$p < 0.0001$.

*Figure S2. Behavioral effects of optogenetic stimulation of NE terminals in BLA*

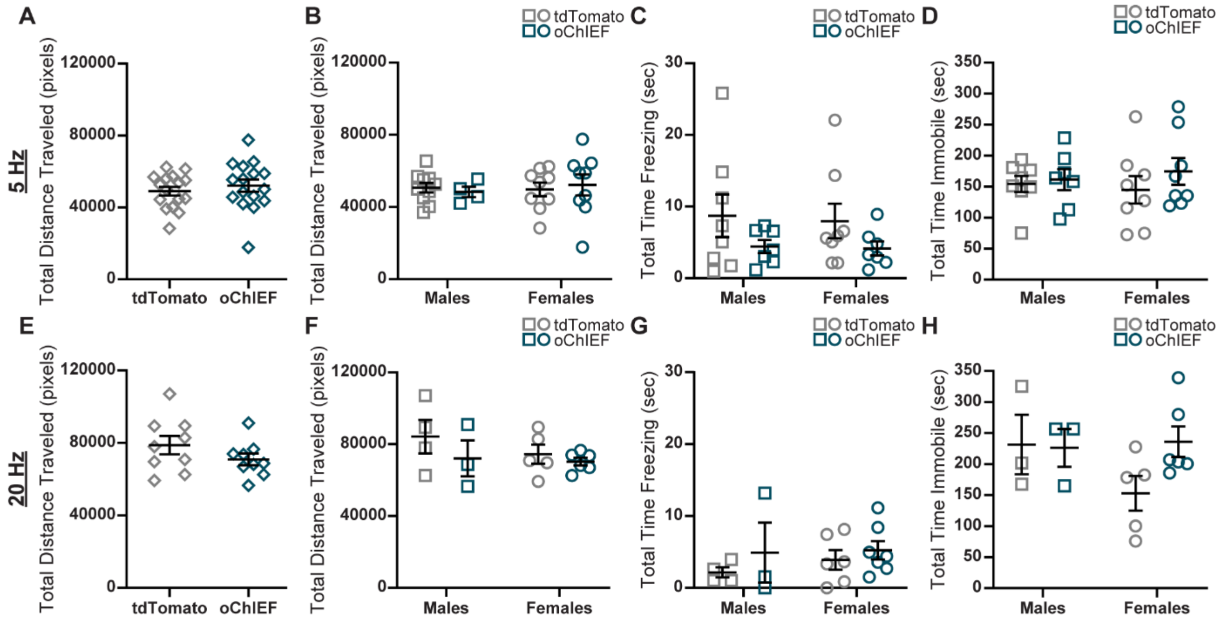


**A.** Effects (mean ± SEM) of 5 Hz stimulation on locomotor activity during OFT ($n$ = 16 mice/group). **B.** Sex-specific effects (mean ± SEM) of 5 Hz stimulation on locomotor activity during OFT ($n$ = 4-10 mice/group), **C.** passive coping during LST ($n$ = 7-8 mice/group), and **D.** passive coping during TST ($n$ = 7-8 mice/group). **E.** Effects (mean ± SEM) of 20 Hz stimulation on locomotor activity during OFT ($n$ = 9 mice/group). **F.** Sex-specific effects (mean ± SEM) of 20 Hz stimulation on locomotor activity during OFT ($n$ = 3-6 mice/group), **G.** passive coping during LST ($n$ = 3-7 mice/group), and **D.** passive coping during TST ($n$ = 3-6 mice/group). Squares indicate males, circles indicate females; gray points indicate tdTomato, teal points indicate oChIEF.

*Figure S3. Sex-specific effects of optogenetic stimulation of BLA NE terminals on cFos levels in BLA and LC*

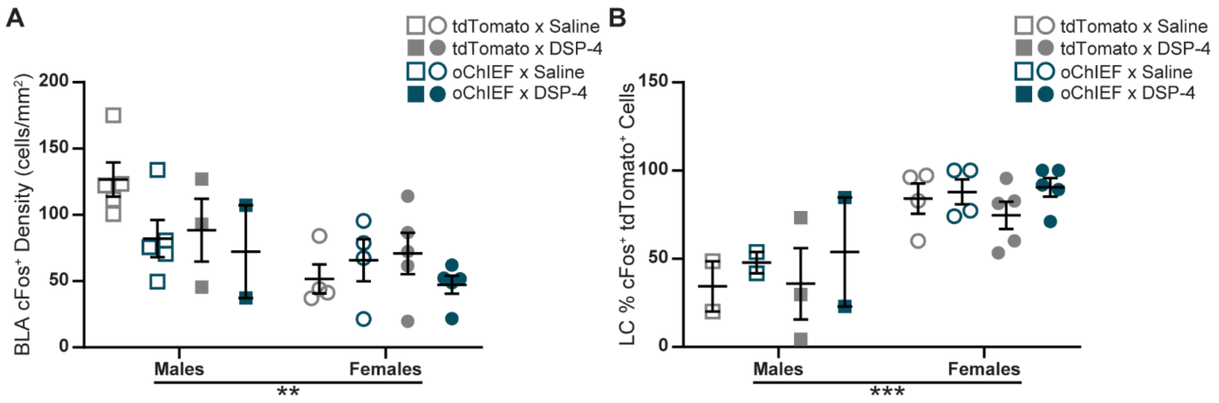


**A.** Effects (mean ± SEM) of DSP-4 administration and 5 Hz stimulation on density of cFos⁺ cells in BLA (*n* = 2-5 mice/group) and **B.** percentage of cFos⁺ cells expressing the viral fluorophore tdTomato in LC (*n* = 2-5 mice/group). Squares indicate males, circles indicate females; open points indicate saline, closed points indicate DSP-4; gray points indicate tdTomato, teal points indicate oChIEF. $**p < 0.01$; $***p < 0.001$.

*Table S1. Statistical analyses of sex differences in Figures S1-3*

| **Analysis** | ***F / t*** | ***p*** |
|---|---|---|
| Repeated Measures 3-way ANOVA: Z-Scored NE Mean | Fig. S1A | |
| Effect of Time | $F_{3.373,53.97}$ = 10.85 | < 0.0001**** |
| Effect of Sex | $F_{1,16}$ = 1.976 | 0.1790 |
| Effect of Coping Response | $F_{1,16}$ = 0.2697 | 0.6106 |
| 2-way ANOVA: Mean Z-Scored NE Detrended | Fig. S1B | |
| Effect of Sex | $F_{1,7}$ = 1.711 | 0.2322 |
| Effect of Coping Response | $F_{1,7}$ = 2.836 | 0.1360 |
| 2-way ANOVA: Max Z-Scored NE Detrended | Fig. S1C | |
| Effect of Sex | $F_{1,6}$ = 5.075 | 0.0652 |
| Effect of Coping Response | $F_{1,6}$ = 13.62 | 0.0102* |
| 2-way ANOVA: Total Distance Traveled | Fig. S2B | |
| Effect of Sex | $F_{1,28}$ = 0.1065 | 0.7466 |
| Effect of Virus | $F_{1,28}$ = 0.0003 | 0.9861 |
| 2-way ANOVA: Total Time Freezing | Fig. S2C | |
| Effect of Sex | $F_{1,26}$ = 0.0551 | 0.8162 |
| Effect of Virus | $F_{1,26}$ = 3.559 | 0.0704 |
| 2-way ANOVA: Total Time Immobile | Fig. S2D | |
| Effect of Sex | $F_{1,27}$ = 0.0097 | 0.9224 |
| Effect of Virus | $F_{1,27}$ = 0.9434 | 0.3400 |
| 2-way ANOVA: Total Distance Traveled | Fig. S2F | |
| Effect of Sex | $F_{1,14}$ = 0.8413 | 0.3746 |
| Effect of Virus | $F_{1,14}$ = 1.651 | 0.2197 |
| 2-way ANOVA: Total Time Freezing | Fig. S2G | |
| Effect of Sex | $F_{1,16}$ = 0.3275 | 0.5751 |
| Effect of Virus | $F_{1,16}$ = 1.281 | 0.2745 |
| 2-way ANOVA: Total Time Immobile | Fig. S2H | |
| Effect of Sex | $F_{1,13}$ = 1.115 | 0.3103 |

| | | |
|---|---|---|
| Effect of Virus | $F_{1,13} = 1.414$ | 0.2556 |
| 3-way ANOVA: BLA cFos$^+$ Density | Fig. S3A | |
| Effect of Sex | $F_{1,25} = 9.057$ | 0.0059** |
| Effect of Drug | $F_{1,25} = 1.125$ | 0.2990 |
| Effect of Virus | $F_{1,25} = 2.482$ | 0.1277 |
| 3-way ANOVA: LC % cFos$^+$ tdTomato$^+$ Cells | Fig. S3B | |
| Effect of Sex | $F_{1,19} = 23.41$ | 0.0001*** |
| Effect of Drug | $F_{1,19} = 0.0004$ | 0.9842 |
| Effect of Virus | $F_{1,19} = 2.246$ | 0.1504 |
| *$p$ < 0.05; **$p$ < 0.01; ***$p$ < 0.001; ****$p$ < 0.0001. | | |
